# A DNA origami fiducial for accurate 3D AFM imaging

**DOI:** 10.1101/2022.11.11.516090

**Authors:** Pauline J. Kolbeck, Mihir Dass, Irina V. Martynenko, Relinde J.A. van Dijk-Moes, Kelly J.H. Brouwer, Alfons van Blaaderen, Willem Vanderlinden, Tim Liedl, Jan Lipfert

**Affiliations:** Department of Physics and Center for NanoScience, LMU Munich, Amalienstrasse 54, 80799 Munich, Germany; Department of Physics and Debye Institute for Nanomaterials Science, Utrecht University, Princetonplein 1, 3584 CC Utrecht, The Netherlands

## Abstract

Atomic force microscopy (AFM) is a powerful technique for imaging molecules, macromolecular complexes, and nanoparticles with nanometer-resolution. However, AFM images are distorted by the shape of the tip used. These distortions can be corrected if the tip shape can be determined by scanning a sample with features sharper than the tip and higher than the object of interest. Here we present a 3D DNA origami structure as fiducial for tip reconstruction and image correction. Our fiducial is stable under a broad range of conditions and has sharp steps at different heights that enable reliable tip reconstruction from as few as ten fiducials. The DNA origami is readily co-deposited with biological and non-biological samples, achieves higher precision for the tip apex than polycrystalline samples, and dramatically improves the accuracy of the lateral dimensions determined from the images. Our fiducial thus enables accurate and precise AFM imaging for a broad range of applications.

---

Atomic force microscopy (AFM) is a powerful technique to visualize nano-to micrometer-scale structures with sub-nanometer resolution^1^. Consequently, AFM imaging is frequently used in a broad range of applications, ranging from solid-state physics, to nanofabrication, photonics, material science, and the life sciences^2-8^. In particular, AFM imaging has provided unprecedented insights into the structure of biological macromolecules and their complexes^9-16^. For the interpretation and modeling of the imaged structures, high-resolution and undistorted AFM images that reflect the true sample dimensions are desirable. However, AFM images are distorted due to the finite size of the AFM tip, resulting in a dilation of image features similar to the convolution of optical images by the point-spread-function of the imaging system^5^. In general, as long as the tip is much sharper than the feature under observation, the measured profile will closely resemble the true shape. Yet, if the sample contains features whose aspect ratio is comparable to that of the tip, distortions due to the finite size of the tip become significant. To correct for the distortions introduced by the tip one can, in principle, estimate the tip geometry and use it to reconstruct the true specimen shape. Unfortunately, the exact shape of most commercial AFM tips is not precisely known. Moreover, the tip shape is variable, even within the same batch of tips, and can also change during the measurement due to wear or contamination of the tip while imaging.

There are several approaches to solve this problem that the shape of the AFM tip is finite and unknown. Villarrubia showed mathematically that the best possible estimate of the tip shape is achieved using a method called ‘blind tip reconstruction’^17^. The approach is based on exploiting features of the AFM image as broadened, inverted replicas of the tip.

The fidelity and quality of this tip reconstruction depend on the calibration sample containing features with similar or greater sharpness than those of the tip. There are commercially available calibration samples, for example polycrystalline or silicon standards with features sharper than the tip^18^ or nanofabricated tip characterizers^19^. Inconveniently, these types of calibration samples must be measured either before or after the actual measurement of interest. In addition, since these types of calibration samples are typically very hard, the shape of the tip is prone to change due to wear when the sample is scanned, which will deteriorate or invalidate the resulting tip reconstruction^20, 21^. In addition, measuring separate calibration samples cannot correct for the effects of changes in tip shape during measurement.

Consequently, it is desirable to use an internal marker, i.e., a reference sample that is co-deposited with the sample of interest. Using an internal reference sample has the advantage that the tip can be characterized during the measurement, which is experimentally convenient and ensures that reference sample and the sample of interest are imaged with identical parameters, since e.g., molecular deformations depend on the AFM imaging mode, the applied force, and the imaging medium. A common internal marker for biological samples is double-stranded DNA^13, 22, 23^, since it is easy to prepare and handle, biocompatible, and well characterized. However, using DNA as a reference sample works only well for samples with a maximum height similar to DNA (1-2 nm depending on the measurement method and force), whereas for higher structures the tip is not sufficiently characterized, since the tip reconstruction requires calibration features of the same height as the sample of interest. Another choice for internal, non-destructive markers like virus particles, e.g. the rod-shaped tobacco mosaic virus (TMV)^24^., or inorganic nanoparticles^25, 26^. However, these are significantly higher than many biologically relevant samples and do not exhibit sharp features, which limits the quality of a blind tip reconstruction.

To overcome these limitations, we present a DNA origami fiducial that provides a 3D reference sample for AFM tip reconstruction with sharply defined steps of different heights and a height profile well matched for use with a broad range of macromolecular complexes (up to 18 nm). The DNA origami technique enables the self-assembly of large numbers of identical nanostructures at the molecular scale^27, 28^, with customized geometry and almost atomistic structural detail^29, 30^. The resulting nanostructures have been shown to be robust and stable in a variety of conditions and are used in a large range of applications^31-38^. In particular, DNA origami structures have been used as molecular rulers^39^ for fluorescence^40, 41^ and super-resolution microscopy^42^. For AFM imaging, a single-layer rectangular sheet DNA structure has been used as a size reference and positioning platform^31, 43, 44^. Yet, the previous structures are not suitable for tip characterization because of their low height and lack of sharply defined features in the z-dimension.

Our DNA origami fiducial combines several characteristics that make it well-suited for blind tip reconstruction: the structural features of DNA origami structures have been characterized with high resolution, its designed structure contains flat faces in the x-y direction and sharp edges in the z direction, creating a four-step staircase from 1-2 to 15-20 nm, well-matched to macromolecular complexes of interest. The fact that it consists of DNA makes it fully biocompatible and enables straightforward surface deposition alongside other biological macromolecules. In addition, we show that our fiducial can be deposited on various surfaces including bare mica, poly-L-lysine (PLL)-coated mica, and aminopropylsilatrane (APS)-coated mica and imaged both dry and in liquid. Taken together, our fiducial enables straightforward AFM image correction for a wide range of nanostructures.

## Design of the DNA origami fiducial structure

We designed the staircase-like nanostructure built of eight layers of parallel helices arranged on a square lattice^30, 45^ (Figure 1a and Supporting Information Figure S1). The designed length (*L1*) of the structure is 200 base pairs (bp), the maximum height (*H4*) is eight helices and the width (*W*) ten helices. For the design purposes, we model DNA helices as a cylinder with a diameter of 2 nm (helix diameter in B-form DNA is 2 nm) and a length of 0.34 nm per bp. Using these parameters and assuming close packing, the approximate size of the designed structure is 68×16×20 nm^3^ (*L1*×*H4*×*W*).

**Figure 1.**
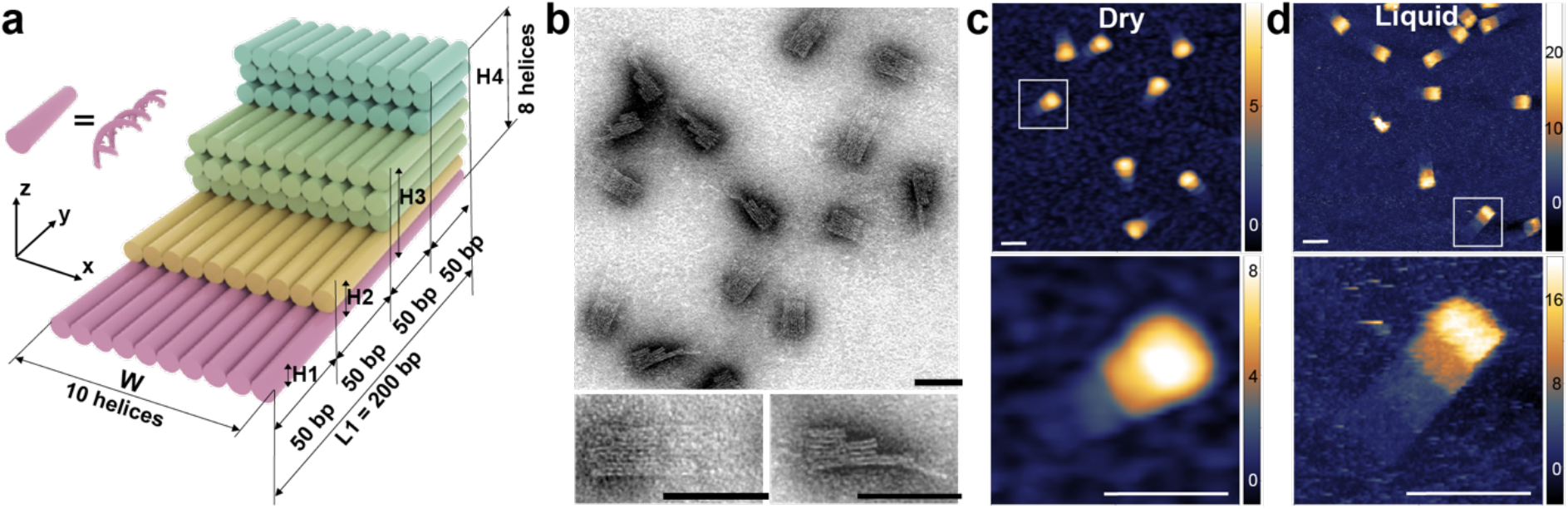
Design of the DNA origami fiducial structure and visualization by TEM and AFM imaging. Design of the 3D DNA origami structure used as a fiducial for AFM imaging. a) Schematic of the designed 3D structure including the design dimensions. The colors represent the different levels. b) Negative-stain TEM images of the fiducial structures confirm the correct assembly and dimensions (a detailed dimension analysis is shown in Supporting Figure S2). The lower images are zoom-ins of two exemplary structures. c) AFM height image of the fiducial structures obtained by imaging on APS mica after drying. The upper and the lower image both have a resolution of 0.5 pixel/nm. The lower image is a zoom-in of the upper image (white box). d) AFM height image of the fiducial structures obtained by imaging on APS mica in liquid. Both images have a resolution of 1 pixel/nm. The lower image is a zoom-in of the upper image (white box). The scale bars in all panels are 50 nm. Z-ranges are indicated by the scale bars on the right.

We varied the number of DNA helices in the layers to obtain four discrete steps with equal areas and heights of one, two, five, and eight helices. Therefore, the fiducial structure features different heights between 2.0 nm and 16 nm which cover a height range suitable for a broad range of samples, including other DNA origami structures^30, 46, 47^ and biological samples^11, 13, 48-50^. Critically, the structure provides sharp and defined vertical 90-degree edges of the steps which is desirable for a reliable AFM tip estimation via blind tip reconstruction.

The *x*-*y* dimension of each step in our design is approximately 17×20 nm (50 bp×10 helices), which provides a sufficient number (>20 points per height plateau) of independent measurement points for calibration^51^ and ensures mechanical stability during the AFM measurement. We used the square lattice geometry and corrected the design for internal twist^45, 52^ to obtain a flat surface of the ‘stairs’.

### Confirmation of correct folding and visualization of the fiducial structures

We folded the fiducial structures in Tris/EDTA/MgCl_2_ buffer, purified excess DNA staple strands, and imaged them with negative-stain transmission electron microscopy (TEM; Figure 1b). The fiducial structures appear as rectangular four-”stair” structures with visible striations running along the length of the fiducial, confirming the direction of the DNA helices. The observed structures lay in different orientations on the surface, while defective or deformed fiducials were not observed, which confirms successful and high-yield assembly of our fiducial structures. Using the TEM images, we analyzed the dimensions of the fiducial structures (Supporting Information Figure S2) and find *L1* = (71.7 ± 3) nm, *H4* = (20.1 ± 1.2) nm, and *W* = (24 ± 1.2) nm, indicating an effective diameter of the DNA helices of (2.51 ± 0.17) nm (Supporting Information Table S1). The effective diameter of DNA helices in 3D DNA origami may vary significantly depending on the DNA origami design, type of packing of the adjacent helices in lattices, number of connecting crossovers and position of nicks^51, 53, 54^. In addition, the spacing of helices depends on solution conditions, like pH, temperature, and in particular ion concentration, as the highly negatively charged helices tend to repel each other electrostatically^55^ resulting in increase of interhelical distance and a “swelling” of the structure with decreasing ionics strength^53, 54^. Our value from TEM analysis is in good agreement with the value determined previously under similar conditions for a multi-layer origami also by TEM (2.25 nm)^51^ and is very close to but slightly smaller than the values determined by SAXS in free solution (∼2.7 nm)^53^ and cryoEM structure modeling (∼ 2.6 nm)^56^.

Next, we imaged the fiducial structures both in situ (i.e., fully hydrated under buffered solution) and after drying in air. In the latter experiment (Figure 1c), we investigated different surface deposition strategies: bare mica, APS mica, and PLL mica. All surface deposition strategies resulted in overall similar images, however, with slight differences in the exact dimensions. (Supporting Information Figure S3). Notably, almost all fiducials were oriented with their large flat face on the substrate’s surface, exposing the staircase feature to the AFM tip, which is also the preferred orientation for our purposes. The heights of the stair steps were almost a factor of 2 lower compared to the heights obtained via TEM (Supporting Information Figure S4 and Table S1), which is expected for dry AFM measurements^57^. Furthermore, it is apparent from the AFM images that the lateral *x-y* dimensions are distorted, in particular the higher features of the staircase appear wider than the lower features (Figure 1c), which is to be expected due to the finite size and conical shape of the tip. In AFM images obtained in liquid (Figure 1d), we see different orientations of the fiducial structure on the surface, however the flat side of the staircase is still attached to the bottom most frequently. Compared to the dry measurements, the structures are significantly higher and appear less distorted, resulting in dimensions close to those measured in the TEM images.

### Fiducial structures enable AFM tip characterization via blind tip reconstruction and subsequent correction for the finite AFM tip size

To test the performance of our DNA origami fiducial structure for estimation of the 3D shape of the AFM tip during measurement, we deposited DNA origami fiducial structures on APS mica and imagined a large field of view (Supporting Information Figure S3b). We then used the features of the DNA origami fiducials to perform blind tip reconstruction following the protocol of Villarrubia^17^ implemented in the image analysis software SPIP (Figure 2a; see Methods in Supporting Information for details). In a next step, we used this tip estimate to correct a zoom-in of the same AFM image scanned with the same tip (Figure 2b,c). To assess the effectiveness of this method, we determined the width of the fiducial structure before and after correction and found that the width was reduced from 31.2 nm to 21.2 nm (Figure 2e), which is much closer to the width of the origami design (Figure 1a) and the width measured in TEM images (Supporting Information Figure S2). To highlight the finite tip size correction, we also calculated a difference image of the corrected image and the original image (Figure 2d) which shows that especially the widths of the higher steps of the staircase were significantly overestimated in the original image.

**Figure 2.**
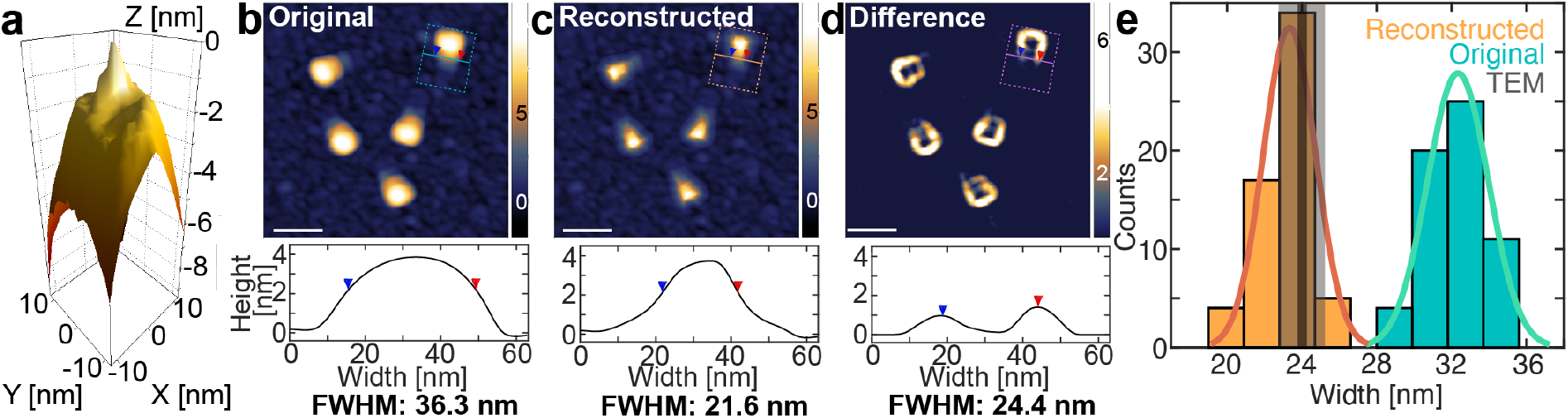
AFM tip characterization and finite tip size correction using the fiducial. a) Estimate of the AFM tip shape obtained by blind reconstruction using the image shown in Supporting Information Figure S3b. b) Top: AFM height image of the fiducial structures on APS mica (dry) with a resolution of 1 pixel/nm. Bottom: height profile of one exemplary molecule, averaged along the fiducial’s long axis (as indicated in the AFM image). Arrows indicate the FWHM^63^; the apparent width is significantly larger than the expected width from the design of ∼22.5 nm and the width measured in negative stain TEM, (23.0 ± 1.2) nm. c) The same image as in panel b after reconstruction based on the tip shape in panel a from blind tip reconstruction. The apparent width now is much closer to what is expected from the design. d) Difference image visualizing the effect of image deconvolution. Scale bars are 50 nm and z-ranges are indicated on the right. e) Width distribution from AFM images before (turquoise) and after (orange) image reconstruction. The solid lines are Gaussian fits. The width of (32.3 ± 1.6) nm (mean ± std) is corrected to (23.3 ± 1.4) nm after correcting for the finite tip size. The corrected value is in excellent agreement with the designed width and the width measured in negative stain TEM indicated by a dark gray vertical line and std in light grey (see Supporting Information Table S1 for a detailed dimension comparison).

### Evaluation of the number of fiducial structures required for reliable tip reconstruction

Having demonstrated that we can estimate the size of the AFM tip by using our reference structures well enough to reproduce the true dimensions of the fiducial structure, we investigate how many fiducial structures are sufficient to get a good estimate of the AFM tip shape. From an image with 114 fiducial structures in total (Figure 3a), we selected between 1 and 100 structures for the blind tip reconstruction. For each number of structures, we randomly selected (with repeats) 20 sets of fiducial structures as inputs for the tip reconstruction. The resulting x and y-profiles of the estimated tip shape (Figure 3b,c; the number of fiducials used increases from blue to red) converge towards the final tip shape result (100 fiducial structures, dark red line) from about ten selected fiducials (Figure 3d). The results suggest that using a minimum of ten fiducial structures for blind tip reconstruction is sufficient for an acceptable tip estimate, in a 1 × 1 μm^2^ image this would for example correspond to a concentration of ∼1 nM.

**Figure 3.**
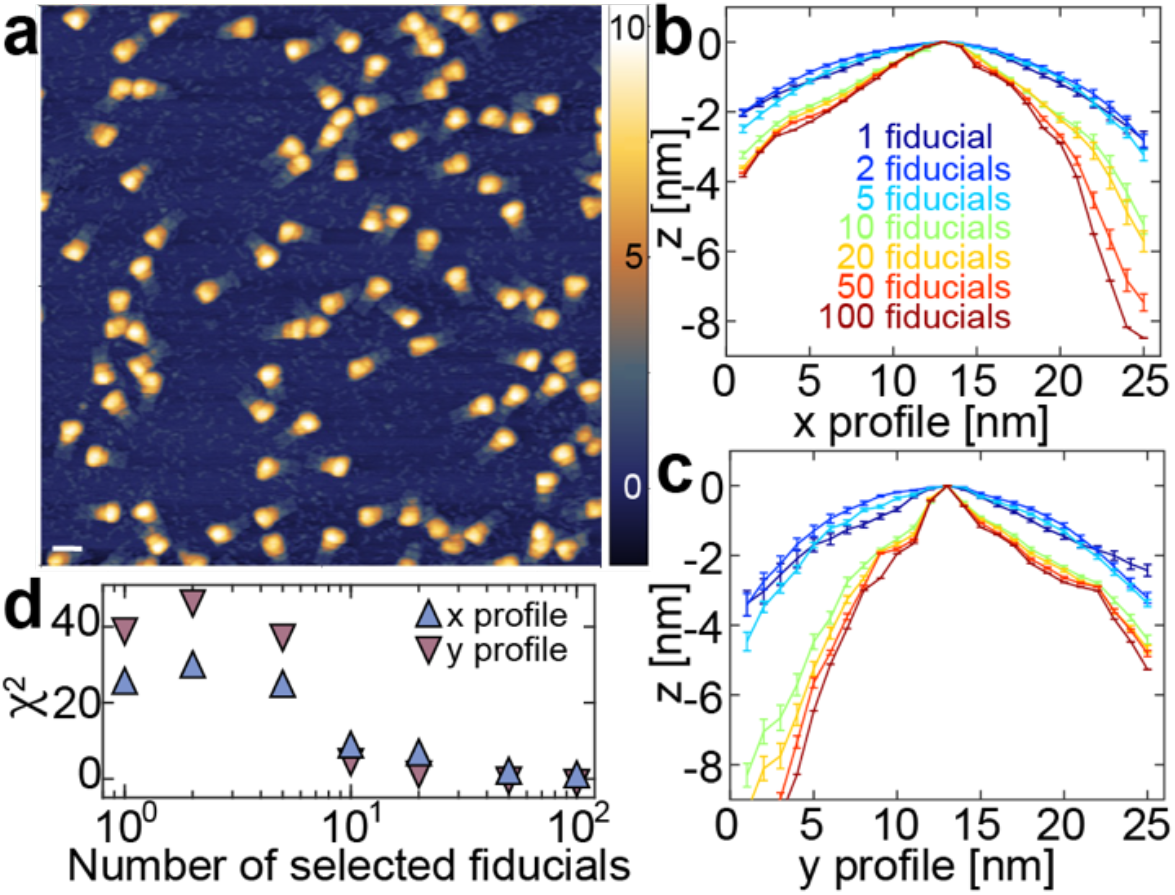
AFM tip characterization using different numbers of fiducials. a) AFM height image of fiducial structures (114 in total) imaged in dry AFM mode on APS mica with a resolution of 1 pixel/nm. The scale bar is 50 nm and the z-range is indicated on the right. b) Estimate of the AFM tip shape x-profile obtained by using between 1 and 100 fiducials for the blind tip reconstruction in the image shown in panel a. For each graph, 20 sets of fiducials were randomly selected, with repeats. c) AFM tip shape y-profile, analogous to panel b. d) χ^2^ of the x- and y-profiles shown in panels b and c as a function of the number of selected fiducials compared to the estimate using 100 fiducials with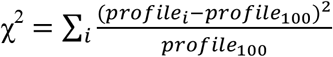.

### Comparison of tip characterization using our fiducial or a polycrystalline sample

Having established an effective method of finite-size tip correction using a DNA origami fiducial structure, we compared our method to correction using an external polycrystalline tip characterization sample (NanoAndMore GmbH, Wetzlar, Germany) with hard sharp pyramidal nanostructures with base length in the range 50 - 100 nm and height 50 - 150 nm, and radius of curvature of the sharpest edges below 5 nm. We characterized five different AFM cantilevers (FASTSCAN-A, Bruker Inc., Billerica, MA, USA; which are used throughout the study) both using our fiducial as well as a polycrystalline sample (Supporting Information Figure S5). Interestingly, we found that for a sharp tip (Supporting Information Figure S5a-c) we get an extremely good estimate of the very edge of the tip when using the DNA fiducial, almost identical to the vendor specifications and significantly better than that the estimate obtained with the polycrystalline sample. The advantage of the polycrystalline sample is that it can characterize a larger z-range of the tip (20-30 nm instead of 5-10 nm for the fiducial sample). For a contaminated or blunt tip (Supporting Information Figure S5d and e), both samples give equally poor results. The results highlight significant tip-to-tip variation even for fresh tips from the same batch. We find that while the polycrystalline sample has the advantage of covering a larger range of heights, for the height range accessible our DNA origami fiducial, it provides higher-resolution tip reconstruction.

### Co-deposition of fiducial structures allows reconstructing the size of a 24-helix bundle DNA origami structure

Given that approximately ten fiducial structures are sufficient to obtain reliable tip shape estimates, we next tested the application of our fiducial *in situ* by co-deposition with another structure of interest. We deposited an equimolar mixture of the fiducial sample and a 24-helix bundle DNA origami (24-HB; Figure 4a). In a 1 × 1 μm^2^ image (1024 × 1024 pixels), we selected 25 fiducial structures for blind tip reconstruction to ensure convergence of the tip estimate (Figure 4b,c). We then used the tip to correct the dimensions of the 24-HBs (Figure 4d-f). The width of the 24-HB is (22.1 ± 1.8) nm (mean ± std; Figure 4f) in the original image, which is significantly larger than the expected width from the design of 15.7 nm^53^. In contrast, in the corrected image, we find a width of (16.3 ± 1.6) nm, which is very close to the value expected from the design. The results suggest that co-deposition of our fiducial provides a convenient and straightforward way to obtain accurate, high-resolution, tip-corrected images.

**Figure 4.**
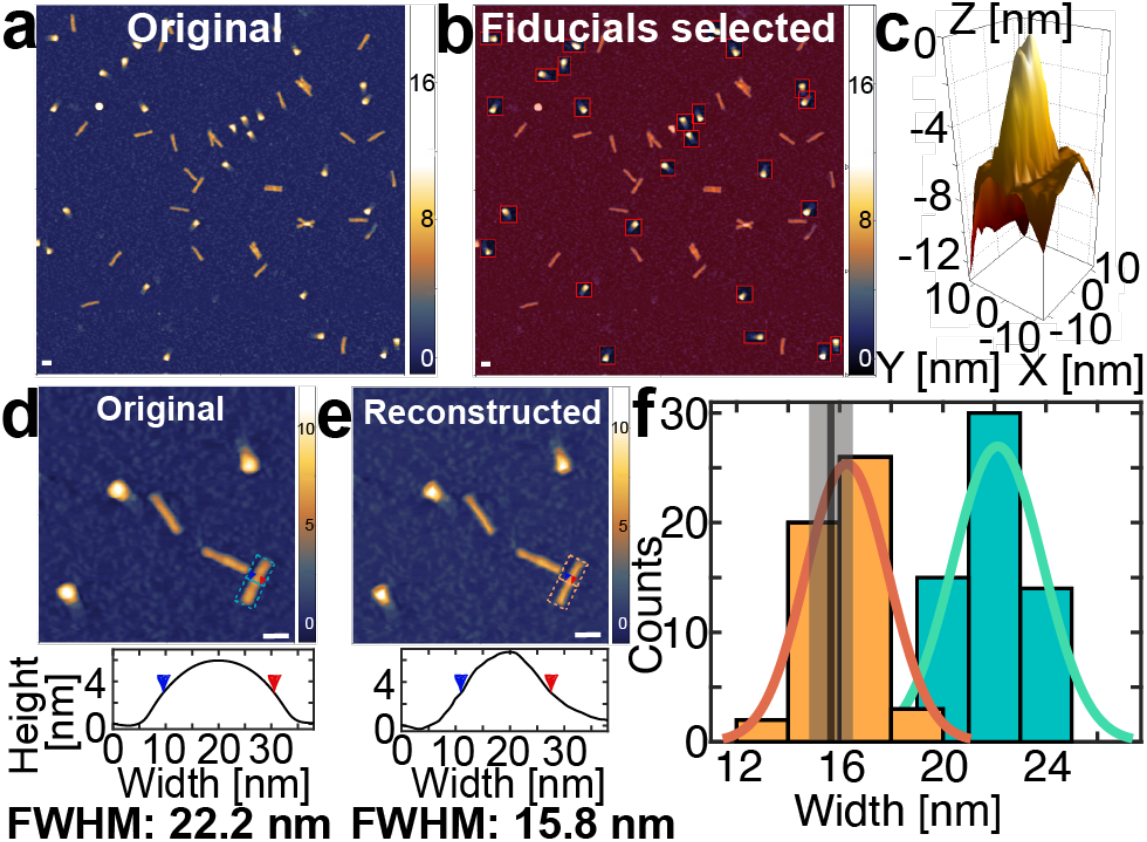
Fiducials enable accurate AFM measurements of a 24-helix bundle DNA origami structure. a) AFM height image of the fiducial structures co-deposited with a DNA origami 24-helix bundle (24-HB) structure, both at a concentration of 1 nM, deposited on APS mica and measured dry with a resolution of 1 pixel/nm. b) Same image as in panel a, with the 25 fiducial structures highlighted by red boxes. The image sections highlighted by the boxes are used for blind tip shape reconstruction. c) Tip shape obtained from blind tip reconstruction using the fiducial samples marked in panel b. d) Top: zoom-in on a different spot of the same sample shown in panel a, the resolution is 2 pixel/nm. Bottom: height profile of the raw AFM image of an exemplary 24-HB. Arrows indicate the FWHM; the apparent width of 22.2 nm is significantly larger than the expected width from the design of ∼15.5 nm. e) Top: the same image as in panel d, after reconstruction based on the tip shape from blind tip reconstruction shown in panel c. All scale bars are 50 nm and Z-ranges are indicated on the right. Bottom: height profile of the reconstructed AFM image of the same 24-HB. f) Width distribution from AFM images before (turquoise) and after (orange) image reconstruction. The solid lines are Gaussian fits. The width of (22.1 ± 1.8) nm (mean ± std) is corrected to (16.3 ± 1.6) nm by finite tip size correction. This value is in excellent agreement with the designed width of (15.5 ± 1.0) nm, indicated by a dark gray vertical line and std in light gray.

### *In situ* image correction for DNA-protein complexes

To demonstrate the applicability of our method to macromolecular complexes beyond DNA origami, we co-deposited our fiducial with double-stranded DNA and a DNA-interacting protein (HIV-1 integrase). We find that the fiducial is bio-compatible and preserves its shape despite the presence of DNA-interacting proteins (Figure 5a). Here again, we observed that the widths of DNA, protein, and protein-DNA complexes are reduced after image reconstruction. As a proof of principle, we compared the DNA width before and after reconstruction and find that it reduces significantly from ∼5.1 nm to ∼2.9 nm (Figure 5b,c), which is much closer to 2 nm, the crystallographic width of double-stranded DNA.

**Figure 5.**
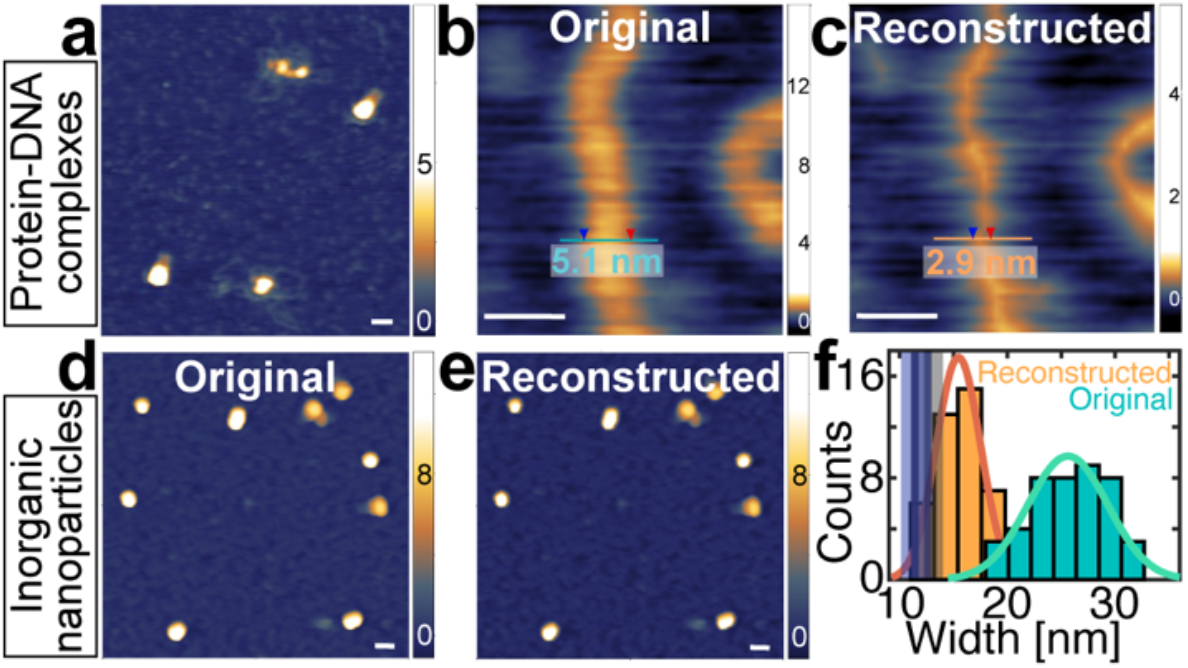
Fiducials enable accurate AFM measurements of DNA protein complexes and inorganic nanoparticles. a) AFM height image of the fiducial structures co-deposited with DNA-protein complexes (DNA length 4.8 kbp; protein HIV-I integrase) deposited on APS mica and measured dry with a resolution of 0.4 pixel/nm. b) Zoom-in on a different spot of the same sample shown in panel d, the resolution is 1.4 pixel/nm. The apparent DNA width of 5.1 nm is significantly larger than the expected DNA width of 2 nm. c) The same image as in panel e, after reconstruction based on the tip shape from blind tip reconstruction as shown in panel d. The DNA width in the reconstructed AFM image is much closer to the expected DNA width of 2 nm. The scale bars are 10 nm and Z-ranges are indicated on the right. d) AFM height image of the fiducial structures co-deposited with SiO_2_ nanoparticles, both at a concentration of 1 nM, deposited on APS mica and measured dry with a resolution of 1 pixel/nm. e) The same image as in panel a, after reconstruction based on the tip shape from blind tip reconstruction using the co-deposited fiducials. The scale bars are 50 nm and Z-ranges are indicated on the right. f) Width distribution from AFM images before (turquoise) and after (orange) image reconstruction. The solid lines are Gaussian fits. The width of (25.6 ± 3.7) nm (mean ± std) is corrected to (15.5 ± 2.1) nm by finite tip size correction. This value is much closer to the imaged height (12.6 ± 1.4) nm (mean ± std), indicated with a dark grey vertical line and the standard deviation in light grey, and also to the width measured in TEM, (11.5 ± 1.2) nm, indicated by a dark blue vertical line and the standard deviation in light blue.

### Height and width analysis of inorganic nanoparticles

Next, we tested our fiducial structure for *in situ* image reconstruction of inorganic nanoparticles. We co-deposited SiO_2_ nanoparticles with our fiducial. Similar as for the biological samples, the apparent widths of the silica nanoparticles are significantly larger than the widths in the corrected image (Figure 5d-f). After image reconstruction, the width is reduced from (25.6 ± 3.7) nm (mean ± std) to (15.5 ± 2.1) nm, which is much closer to the width measured in TEM images (11.5 ± 1.2) nm (Supporting Information Figure S6a,b) and also to the height of the particles measured in AFM, (12.6 ± 1.4) nm (Supporting Information Figure S6c,d). After image correction, the AFM-determined width and height measurements and the TEM-derived widths are in overall approximate agreement, as would be expected for spherical particles. The fact that the dimensions from the TEM analysis are still slightly smaller than AFM derived values might be due to the fact the ultra-high vacuum used during TEM imaging leads to a small reduction in particle size^58^. The remaining small difference between AFM-determined width and height might stem from imperfections in the image correction, from slight compression of the particle by the AFM cantilever, or could be due to the fact that particles are not perfectly spherical and tend to adhere to the surface with their larger side. Additionally, we observed that the width distribution becomes smaller after image reconstruction (Figure 5f) while the height distribution stayed the same within error (Supporting Information Figure S6d). The reduction in the variance of the width distribution is likely due to an asymmetry of the tip, such that the widths of the particles are distributed wider in the original image than in the reconstructed image.

In conclusion, we have established a method that allows us to correct for the finite size of the AFM tip and its specific shape while scanning a sample, employing a DNA origami staircase structure as a fiducial for AFM image calibration. We demonstrate that our fiducial structures are versatile and biocompatible, can be deposited on various surfaces including bare mica, APS mica, and PLL mica, and can be imaged in liquid or dry. This allows straightforward surface co-deposition with samples of interest, and we demonstrate the broad applicability of the method by imaging DNA origami structures, DNA-protein complexes, and silica nanoparticles. In all cases, the blind tip reconstruction using our fiducial allows for subsequent correction of images for finite tip size, which enables much more accurate determination of sample dimensions than uncorrected images. We show that as few as ten fiducial structures are sufficient for a good tip estimate. Also, we demonstrate that to characterize the very edge of the tip of FASTSCAN-A cantilevers, the fiducial sample gives a better estimate of the tip shape than a commercial polycrystalline characterization sample. Taken together, our method enables accurate, straightforward, and user-friendly AFM image correction.

We anticipate many new applications coming within reach by using DNA origami structures as fiducials for 3D AFM image calibration. We note that the design of the DNA origami structure could be altered or extended for specific purposes, for example by addition of additional steps or attachment of fluorescent dyes. A combination with fluorescent markers has the potential to enable simultaneous use of the fiducial as a tip shape and fluorescence calibration^39-42, 58, 59^. Another research direction would be to go beyond imaging and to use the fiducial as a mechanical stiffness marker to study the compliance of biomolecules to indentation forces^60^, e.g. to probe the effects of silicifaction^61, 62^ or other functionalization. We, therefore, anticipate that our fiducial will provide a multi-modal calibration platform for a range of applications.

## Author Contributions

P.J.K. and W.V. designed this study. M.D. and I.M. designed, assembled, and purified DNA origami samples; R.J.A.v.D-M., K.J.H.B., and A.v.B. synthesized and characterized silica nanoparticles; P.J.K. and W.V. performed AFM measurements and analyzed the data. T.L. and J.L. supervised research; P.J.K. and J.L. wrote the manuscript with input from all authors.

## Notes

The authors declare no competing financial interest.

## Acknowledgements

We thank Thomas Nicolaus for laboratory assistance, Diogo Saraiva, Lisa Tran, Steven De Feyter and Aidin Lak for helpful discussions, and Arthur Ermatov for critical reading of the manuscript.

