## Supplementary Information for "A DNA origami fiducial for accurate 3D AFM imaging"

**Supplementary Materials and Methods**
**Supplementary Table S1**
**Supplementary Figures S1-S6**

**Supplementary Materials and Methods**

**DNA origami design and assembly**

The DNA origami AFM fiducial structure was designed using caDNAno[1] (design schematics in Figure S1). The staircase-like structure consists of eight layers of parallel helices packed on a square lattice. The designed length of the structure is 200 base pairs, the width is ten helices. The number of DNA helices in the layers is varied to obtain four discrete steps with equal x-y areas and heights of one, two, five, and eight helices.

Design-specific staple strands were purchased from IDT Technologies, the scaffold strand p8634 was produced from M13 phage replication in E. coli. Scaffold strand and staple strands were mixed at 1:5 scaffold:staple ratio with target concentrations of 30 nM and 150 nM (each staple), respectively in 10 mM Tris Base, 1 mM EDTA buffer with 18 mM magnesium chloride (TE/Mg2+). 50 µL volumes of staple/scaffold mixture were heated up to 65◦C for 5 min and annealed from 65◦C to 20 °C at -0.2°C/min in a PCR machine. The DNA origami structures were purified from excess staples using 100 kD molecular weight cut-off filters (Amicon Ultra-0.5 Centrifugal Filter Units with Ultracel-100 membranes). The 24-helix-bundle (24HB) structure [ 10.1021/acs.nanolett.6b01335] was folded in a similar fashion using the p8064 scaffold strand and purified using the PEG precipitation method adapted from [10.1002/cbic.201700377].

**TEM sample preparation and imaging**

5 µL of sample solution was incubated for 30 s – 5 min, depending on concentration, on glow discharged TEM grids (formvar/carbon, 300 mesh Cu; Ted Pella) at room temperature. After incubation on the grids, the sample was wicked off by bringing the grid into contact with a filter paper strip. Samples containing DNA origami went through an additional staining step with a 2% uranyl formate aqueous solution containing 25 mM sodium hydroxide. After incubating and wicking the sample off, a 5 µL drop of staining solution (2% uranyl formate aqueous solution containing 25 mM sodium hydroxide) was applied to the grid, immediately wicked off, followed by applying another 5 µL drop of staining solution. This drop was allowed to incubate on the grid for 10 seconds and then wicked off. The grid was allowed to dry for 5 minutes before imaging. Imaging was performed with a JEM1011 transmission electron microscope (JEOL) operated at 80 kV.

**Synthesis of $SiO_2$ nanoparticles**

Silica particles were synthesized with a one-step synthesis based on previous literature[2, 3]. All glassware was etched of residual silica via a base bath (2−3 days in a saturated solution of KOH in isopropanol, rinsed with milliQ water). The particles were synthesized as follows: in a 500 mL 1-neck flask, 181 mg (6 mM) L-arginine (98%, Sigma-Aldrich) were dissolved in 169 mL milliQ water. The mixture was heated to 30 °C and stirred slowly (200 rpm). After 1 h, 11.2 mL (49 mmol) TEOS (tetraethoxysilane; 98%, Sigma-Aldrich) was added slowly via the wall and a two layered system formed (top: TEOS, bottom: water). The reaction mixture was stirred for 1 week to complete the synthesis. The resulting particles were stored at room temperature in the dark and used without further processing.

**AFM sample preparation**

For the AFM samples, we deposited 20 µL of a buffered solution (10 mM Tris Base, 12.5 mM $MgCl_2$,1 mM EDTA, pH 8.35; AFM buffer) containing the fiducial structures (at different concentrations between 1 and 10 nM) either on freshly cleaved bare muscovite mica or on amino-propyl-silane (APS)-coated mica or poly-L-lysine coated mica. The sample was incubated 5 minutes before washing with 20 mL milliQ water and drying with a gentle stream of filtered argon gas. The APS coating was performed following the protocol from Shlyakhtenko et al.[4]. The poly-L-lysine coated mica was prepared as described previously[5] by depositing 20 µL 0.01%-poly-L-lysine on freshly cleaved muscovite mica for 30 seconds and subsequently rinsing the surface with 30 mL of milliQ water before drying with a gentle stream of filtered argon gas.

For the liquid measurements, 2.5 mL of the buffered solution was added to the sample after incubation. For the co-deposited samples, we pre-mixed the fiducial structures with the corresponding sample (at varying concentrations between 1 and 10 nM) prior to deposition in AFM buffer. The samples were incubated, washed, and dried as described above.

For the DNA-protein complex sample, we first mixed linearized plasmid pU3U5 (4.751 kbp; Mini-HIV DNA, see Cherepanov *et al.*[6]) with HIV-I IN in sodium buffer (10 mM Tris-HCl, 90 mM NaCl; 5 mM $MgCl_2$) to a final concentration of 1 ng/ul DNA and 1 µM of protein. Next, we added the fiducial structures at a final concentration of 1 nM and deposited 20 µL of the mixture on APS-coated mica. The sample was incubated, washed, and dried as described above.

**AFM imaging**

The dry AFM images were recorded in tapping mode at room temperature using the Nanowizard Ultraspeed 2 (JPK, Berlin, Germany) with silicon tips (FASTSCAN-A, drive frequency 1400 kHz, tip radius 5 nm, Bruker, Billerica, Massachusetts, USA). Images were scanned over different fields of view and with various pixel sizes (indicated for each image) with a scanning speed of 5 Hz. The free amplitude varied from 20 to 30 nm. The amplitude setpoint was set to 80% of the free amplitude and adjusted to maintain good image resolution.

The liquid AFM images were recorded in peak-force tapping mode at room temperature also using the Nanowizard Ultraspeed 2 (JPK, Berlin, Germany) with silicon tips (BL-AC40TS, drive frequency 25 kHz in water, tip radius 10 nm, Olympus, Tokyo, Japan). Images were scanned over different fields of view and with various pixel sizes (indicated for each image). The peak force was set to 200 pN.

**AFM image analysis**

Postprocessing of AFM data was performed in the software SPIP (v.6.4, Image Metrology, Hørsholm, Denmark), which has implemented blind peak reconstruction as well as image deconvolution following Villarrubia's protocol[7].

First, the images were flattened and line wise leveled. Next, the tip was characterized via blind tip reconstruction. To this end, we either used the entire image or (in case of co-deposition or contamination of the sample) selected a subset of fiducial structures. Next, we used the tip characterization tool and specified the tip size in x and y as number of pixels. For FASTSCAN-A cantilevers, we took the manufacturer’s specified tip radius of 12 nm as a starting point for the blind tip reconstruction. The resulting tip shape was saved and then loaded to deconvolute the same or another image scanned by the same tip. Here, too, the tip size had to be adjusted to the corresponding size in pixel so that the resolution was not lost.

We note that while we used SPIP for all image processing, other AFM post-processing softwares, such as Gwyddion, have also incorporated blind tip reconstruction routines with implementations very similar to SPIP.

**Supplementary Table**

| Parameter | Design | TEM | Dry AFM | Liquid AFM |
|---|---|---|---|---|
| **Width W** | 10 helices | 24 ± 1.2 nm | 32.3 ± 1.6 nm (original)<br>23.3 ± 1.4 nm (reconstr.) | 30.0 ± 2.2 nm (original)<br>28.8 ± 2.9 nm (reconstr.) |
| **Height H1** | 1 helix | - | 0.65 ± 0.3 nm | 0.55 ± 0.4 nm |
| **Height H2** | 2 helices | 5.1 ± 0.4 nm | 2.1 ± 0.4 nm | 2.0 ± 0.5 nm |
| **Height H3** | 5 helices | 12.8 ± 1 nm | 5.4 ± 0.4 nm | 9.4 ± 1.4 nm |
| **Height H4** | 8 helices | 20.1 ± 1.2 nm | 8.0 ± 0.4 nm | 15.9 ± 1.1 nm |
| **Interhelical spacing (vertical)** | | 2.51 ± 0.17 nm | 1.1 ± 0.2 nm | 2.0 ± 0.2 nm |

**Table S1. Dimension analysis of the DNA origami fiducial structure**
Comparison of the design dimensions to the dimensions measured in TEM, dry AFM, and liquid AFM. Not all features were consistently visible in the different techniques and are therefore not listed. For the TEM data, the mean and standard deviation are listed. Details about the analysis and the raw data can be found in Supporting Information Figure S1. For the dry and liquid AFM data, Gaussians are fitted to the data (Supporting Information Figure S3) and here the mean and sigma of the distribution are listed.

### Supplementary Figures

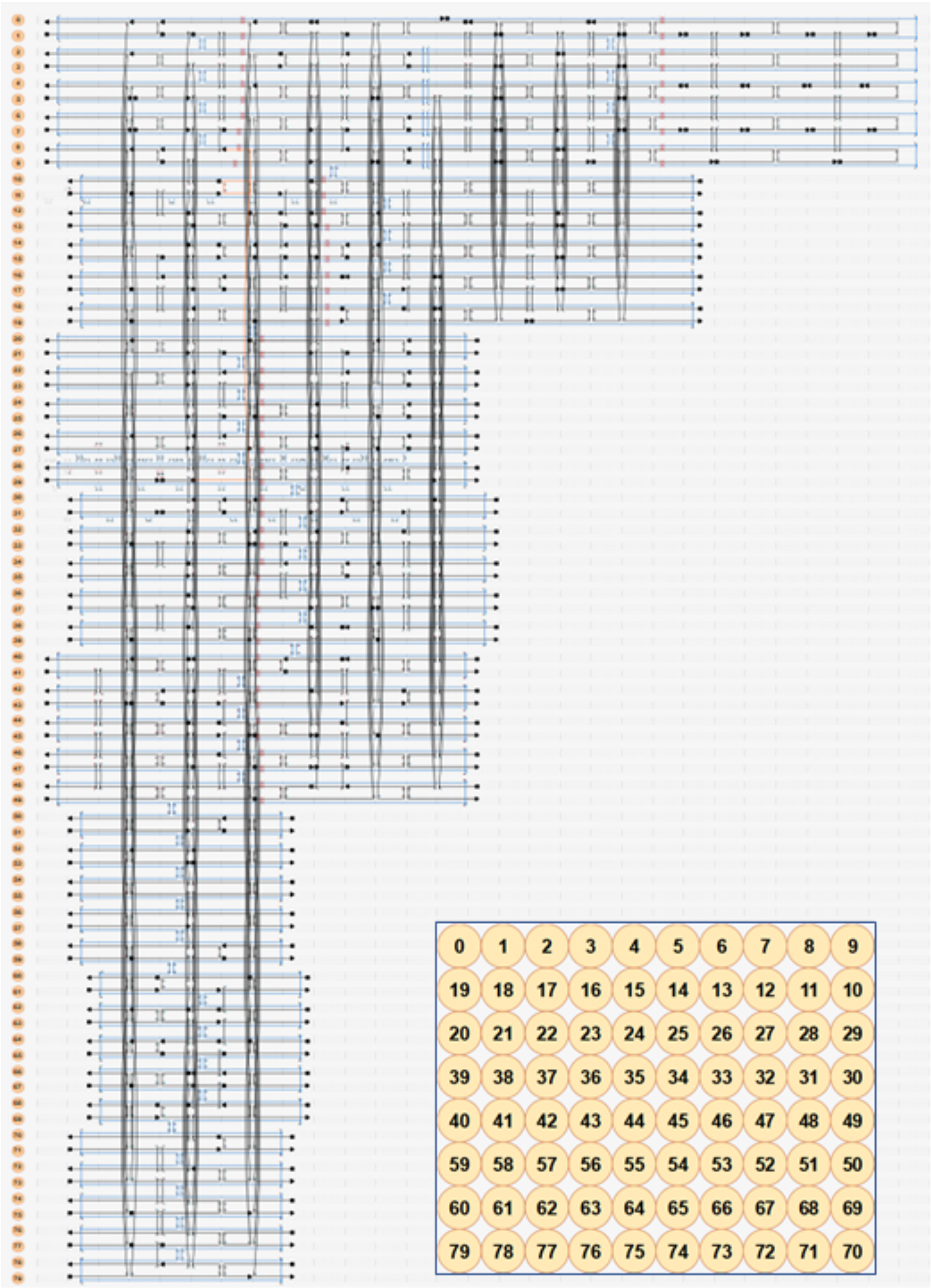


**Figure S1. CaDNAno layout of the DNA origami AFM fiducial structure design.**

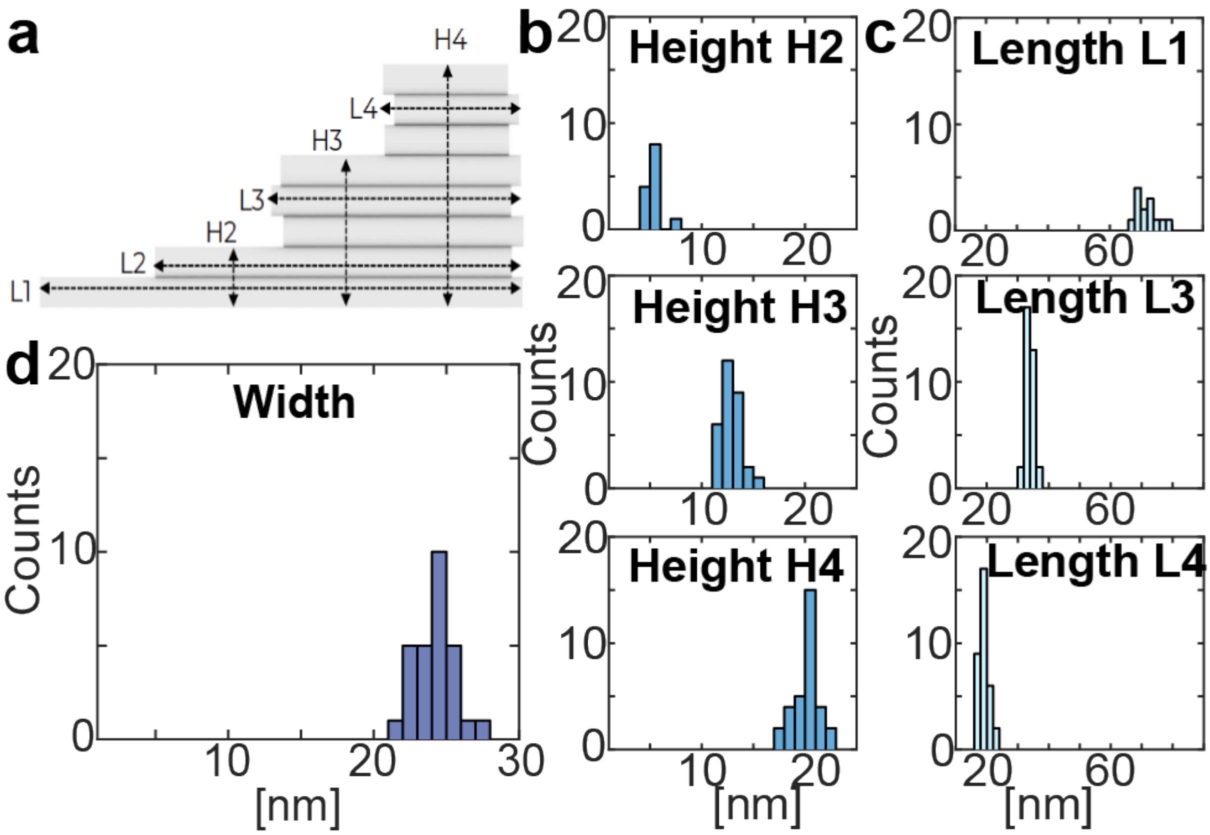


**Figure S2. Dimension analysis of the fiducial structures based on TEM images.** a) Design of the fiducial structure indicating the design dimensions and labelling of the lengths and heights. b) Height distribution for the 3 highest levels of the fiducial (the lowest level H1 was not visible in the TEM images; see Table S1 for a detailed dimension comparison). c) Length distribution for the levels 1, 3, and 4 of the fiducial (length L2 was not clearly visible in the TEM images). d) Width distribution of the fiducial.

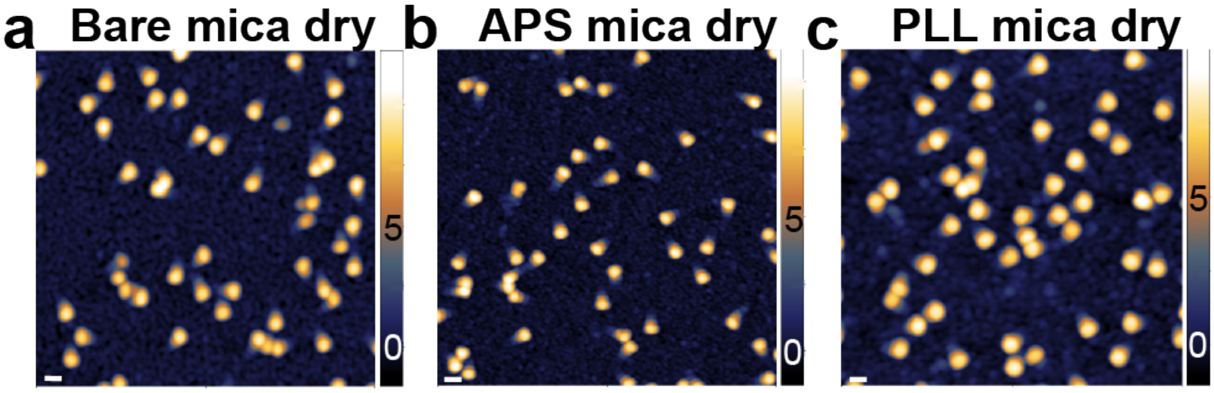


**Figure S3. Comparison of AFM images obtained with different surfaces deposition approaches.** a) AFM height image of fiducial structures at a concentration of 10 nM deposited on a) Mg mica, b) APS mica, c) PLL mica. The scale bars are 50 nm. Z-ranges are indicated by the scale bars on the right.

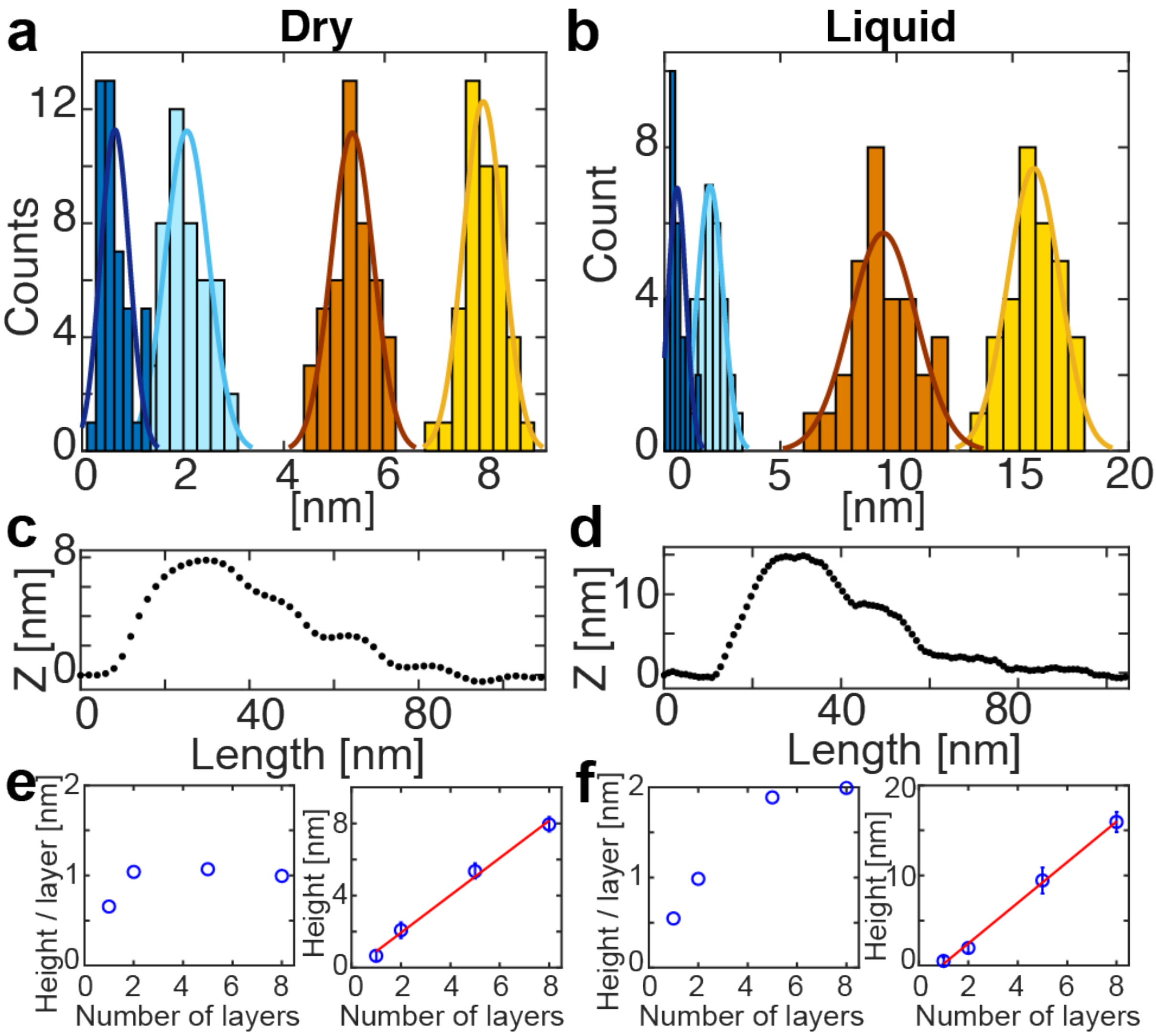


**Figure S4. Dimension analysis of the fiducial structures based on AFM images.** a) Height distribution for the 4 different levels of the fiducial from dry AFM images and Gaussian fits (solid lines) with Gaussians. b) Height distribution for the 4 different levels of the fiducial from liquid AFM images and Gaussian fits (solid lines). See Table S1 for a detailed dimension comparison. c) One exemplary fiducial height profile from dry AFM imaging. d) One exemplary fiducial height profile from liquid AFM imaging. e) Height per DNA layer and total height as a function of the number of DNA layers in the DNA origami for dry AFM imaging. f) Height per DNA layer and global height as a function of the number of DNA layers in the DNA origami for liquid AFM imaging. The red lines in e and f indicate linear fits.

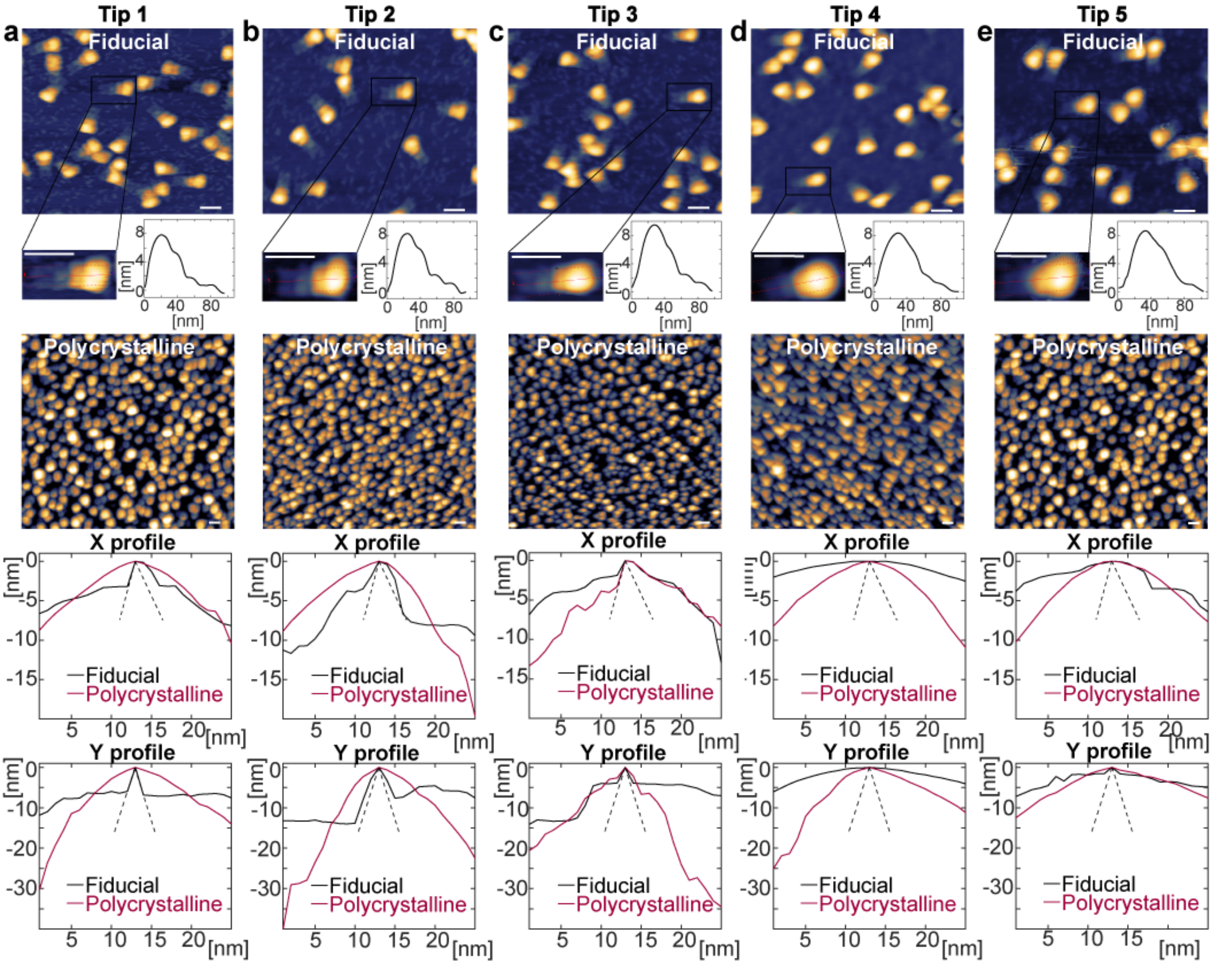


**Figure S5. AFM tip characterization using different AFM FASTSCAN-A cantilevers.** a) Top: AFM height image of fiducial structures at a concentration of 5 nM on APS mica, measured dry. Total image size is 500 x 500 nm$^2$ and 512 x 512 pixels. One exemplary fiducial structure is shown as a zoom-in as well as its height profile. Center: scan of a polycrystalline sample with the same tip. Total image size is 1 x 1 µm$^2$ and 1024 x 1024 pixels. Bottom row: AFM tip shape (height profile along x- and y) obtained from blind tip reconstruction using the fiducial sample or the polycrystalline sample, respectively. As a reference, the tip shape stated by the vendor is co-plotted as a dashed line. b) – e) Analogous to panel a for different FASTSCAN-A tips from the same batch. The data suggest considerable variation from between tips; Tips used for panels a and b enabled high-resolution images and reconstructed tip shapes using our fiducial standard are close to vendor specifications. Tips used for panels c-e appeared less sharp and gave only lower resolution images

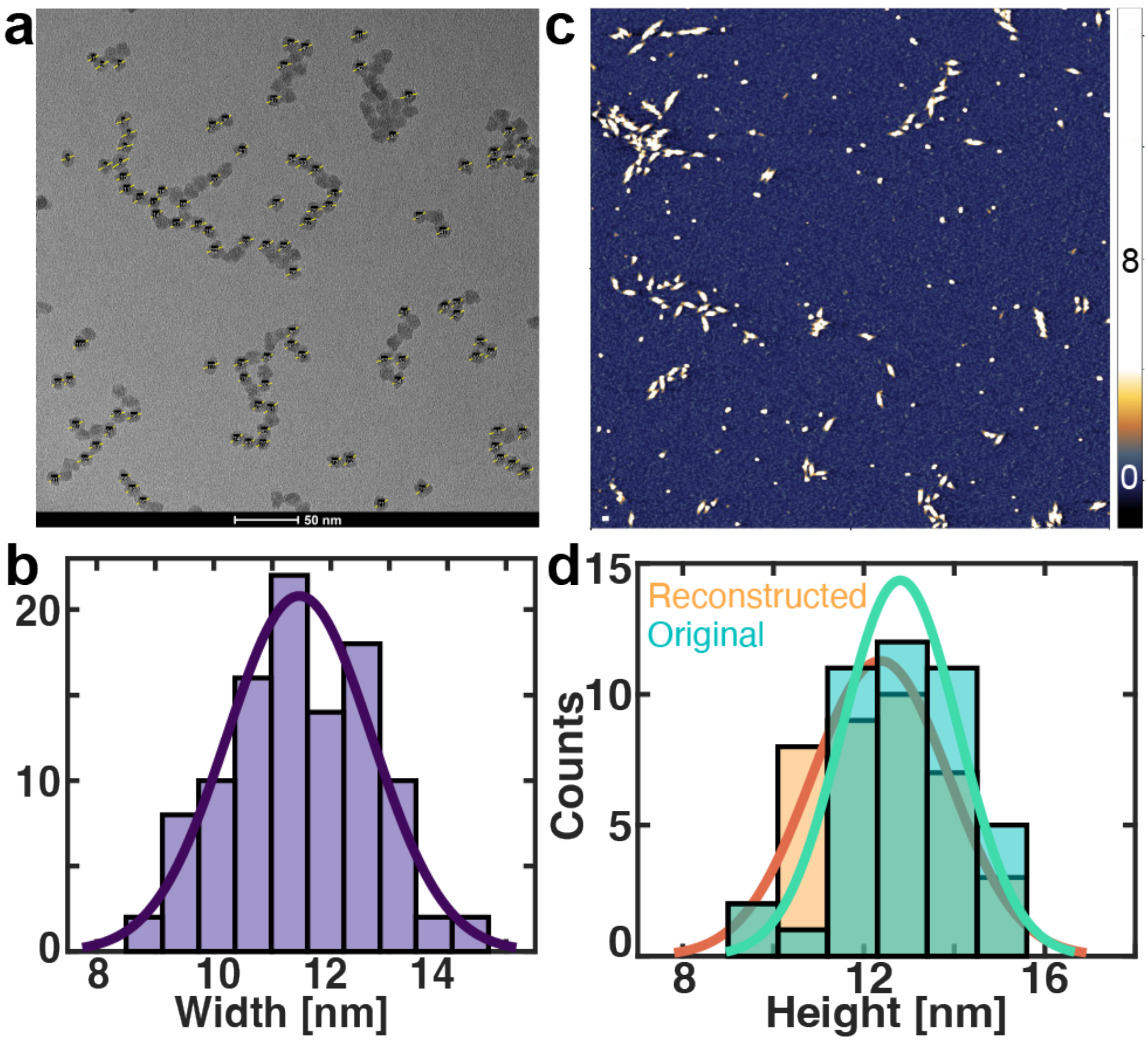


**Figure S6. Width analysis of $SiO_2$ nanoparticles based on TEM images and co-deposition with fiducial structures in AFM with height analysis.** a) TEM image of $SiO_2$ nanoparticles to determine size and shape. Yellow lines indicate cross-sections used for size analysis. b) Width distribution of the $SiO_2$ nanoparticles from the TEM image shown in panel a with a mean and standard deviation of (11.5 ± 1.2) nm. c) AFM height image of the fiducial structures co-deposited with SiO2 nanoparticles, both at a concentration of 1 nM, deposited on APS mica and measured dry with a resolution of 1 pixel/nm. The scale bar is 50 nm. The Z-range is indicated by the scale bar on the right. d) Height distribution from AFM images before (turquoise) and after (orange) image reconstruction. The solid lines are Gaussian fits. The mean height in the original image of (12.8 ± 1.3) nm (mean ± std) does not change within error after image reconstruction (12.4 ± 1.5) nm by finite tip size correction.
